# Multi-scale modelling of location- and frequency-dependent synaptic plasticity induced by transcranial magnetic stimulation in the dendrites of pyramidal neurons

**DOI:** 10.1101/2024.07.03.601851

**Authors:** Nicholas Hananeia, Christian Ebner, Christos Galanis, Hermann Cuntz, Alexander Opitz, Andreas Vlachos, Peter Jedlicka

**Affiliations:** Computer-Based Modelling in the field of 3R Animal Protection, Faculty of Medicine, Justus Liebig University Giessen, Giessen, Germany; Translational Neuroscience Network Giessen, Germany; Charité · NeuroCure (NCRC), Charité Universitätsmedizin Berlin; Department of Neuroanatomy, Institute of Anatomy and Cell Biology, Faculty of Medicine, University of Freiburg, Freiburg, Germany; Dept of Biomedical Engineering, University of Minnesota, Minneapolis, MN, USA; BrainLinks-BrainTools Center, University of Freiburg; Bernstein Center Freiburg, University of Freiburg; Center for Basics in Neuromodulation (NeuroModulBasics), Faculty of Medicine, University of Freiburg, Freiburg, Germany; Ernst Strü ngmann Institute (ESI) for Neuroscience in cooperation with the Max Planck, Society, Frankfurt am Main, Germany; Frankfurt Institute for Advanced Studies, Frankfurt am Main, Germany

**Keywords:** Multi-scale modelling, Dendrites, Magnetic field, Dendritic spike, Backpropagating action potential, STDP, Theta-burst stimulation, Distal/proximal synapses

## Abstract

**Background:** Repetitive transcranial magnetic stimulation (rTMS) induces long-term changes of synapses, but the mechanisms behind these modifications are not fully understood. Although there has been progress in the development of multi-scale modeling tools, no comprehensive module for simulating rTMS-induced synaptic plasticity in biophysically realistic neurons exists..

**Objective:** We developed a modelling framework that allows the replication and detailed prediction of long-term changes of excitatory synapses in neurons stimulated by rTMS.

**Methods:** We implemented a voltage-dependent plasticity model that has been previously established for simulating frequency-, time-, and compartment-dependent spatio-temporal changes of excitatory synapses in neuronal dendrites. The plasticity model can be incorporated into biophysical neuronal models and coupled to electrical field simulations.

**Results:** We show that the plasticity modelling framework replicates long-term potentiation (LTP)-like plasticity in hippocampal CA1 pyramidal cells evoked by 10-Hz repetitive magnetic stimulation (rMS). This plasticity was strongly distance dependent and concentrated at the proximal synapses of the neuron. We predicted a decrease in the plasticity amplitude for 5 Hz and 1 Hz protocols with decreasing frequency. Finally, we successfully modelled plasticity in distal synapses upon local electrical theta-burst stimulation (TBS) and predicted proximal and distal plasticity for rMS TBS. Notably, the rMS TBS-evoked synaptic plasticity exhibited robust facilitation by dendritic spikes and low sensitivity to inhibitory suppression.

**Conclusion:** The plasticity modelling framework enables precise simulations of LTP-like cellular effects with high spatio-temporal resolution, enhancing the efficiency of parameter screening and the development of plasticity-inducing rTMS protocols.

**Highlights:**

- First rigorously validated model of TMS-induced long-term synaptic plasticity in extended neuronal dendrites that goes beyond point-neuron and mean-field modelling
- Robust simulations of experimental data on LTP-like plasticity in the proximal dendrites of CA1 hippocampal pyramidal cells evoked by 10 Hz repetitive magnetic stimulation (rMS)
- Replication of distal synaptic plasticity for a local electrical theta burst stimulation (TBS) protocol
- Prediction of distal and proximal LTP-like plasticity for rMS TBS
- 1 Hz rMS does not induce long-term depression

## Introduction

Transcranial magnetic stimulation (TMS, [1]) is a widely used brain stimulation method which has shown great promise for the treatment of a variety of neurological conditions [2], including stroke [3], pain and drug-resistant depression [4–6]. Further it is also investigated for the use in tinnitus, epilepsy, Alzheimer’s disease [7], Parkinson’s disease [8], schizophrenia [9] and other pathologies. Despite its widespread use, there remain significant gaps in scientific understanding about its mechanisms [10, 11]. As TMS becomes more widespread in both clinical and research applications, it is important to gain a better understanding of the effects of the stimulation at the whole-brain, network, and cellular levels [12, 13].

TMS induces a strong electric field in the brain beneath the TMS coil. This electric field can stimulate whole areas of the brain, and will elicit action potentials most likely in the axonal arbor of affected brain cells [13, 14]. The clinical effects of TMS are related to changes in neuroplasticity [15]. However, the links between TMS-induced electric fields, neuronal spiking and long-term synaptic plasticity are not fully understood.

TMS has been shown to have a variety of effects on the target area. In the CA1 area alone, TMS has been found to increase intrinsic excitability [16], and to induce or modulate LTP [17–26] (see also [27–29]). Importantly, TMS affects neural tissue through electric fields across the whole neuron, which differs from classical LTP induction protocols based on focal direct electric stimulation [28]. To facilitate realistic modelling of LTP induction by TMS, we have used a previously established model of synaptic plasticity [30]. This model is particularly suitable for simulating rMS-induced changes of synapses at neuronal dendrites because it can generate detailed spatio-temporal profiles of synaptic plasticity induced by local compartment-specific voltage threshold-crossing events in the cell such as dendritic spikes, NMDA spikes, large EPSPs, backpropagating somatic spikes, dendritic plateau potentials or combinations thereof [30].

The induction of long-term synaptic potentiation (LTP) by magnetic stimulation has been studied experimentally [31] [32], where magnetic stimulation at 10 Hz induced LTP. This LTP was preferentially located in the proximal dendrites of the cell, with the magnitude of LTP decreasing sharply with further distance from the soma. As there is no cranium in this case, we will henceforth refer to this stimulation as repetitive magnetic stimulation (rMS).

We found that our model, was able to reproduce and better explain the distance-dependent induction of LTP by rMS observed by Lenz et al. [31], as well as to predict a dependence of LTP amplitude on rMS frequency and the ability of rMS to generate LTP in the distal tuft synapses.

## Materials and methods

### Simulation software and code availability

The code will be made available online. For reviewers, a Dropbox link is available here: https://www.dropbox.com/scl/fo/jk4zcgw3c2jefxj23pu3m/ACoeQcrb6DitMmfDS1WDpb rlkey=jjy8b4rvq2z0xk4d4miu0ohhz&st=pylda2tb&dl=0

All *NEURON* simulations were performed using the 7.8.2 release build of *NEURON* [33]. *NeMo-TMS* [34] was run on a modified version of the latest release build (2.0) on *MATLAB R2019b*. *HippoUnit* was run using *Python 3.9*. All simulations and validations were performed on a Windows 10 PC with an Intel Core i9-9900k processor running at 4.7 GHz and 48 GB of RAM.

Since spatial effects are not insignificant in the simulation of magnetic stimulation, simplified point neurons are not appropriate. On the other hand, morphologically fully detailed models are computationally expensive for tuning and running long-term plasticity simulations [35].

Therefore, we implemented our previously established biophysically detailed but morphologically reduced morphology model of the CA1 pyramidal cell [36]. This model features branching dendrites in a simplified but anatomically inspired morphology, and has been validated through the *HippoUnit* test battery [37] including nonlinear dendritic integration and back-propagation of action potentials [36].

For each simulation set, we generated a set of ten models to show the spread of simulation data. To do this, we randomise the synapse locations while keeping the total synaptic input the same. The same set of ten models was used for each simulation set to enable direct comparison.

### Computational model of CA1 pyramidal cell

For all simulations here, we implemented a modified form of the biophysically complex but morphologically reduced model [36] of a CA1 pyramidal cell. The axon was modified with an axon initial segment and an unmyelinated terminal segment as well as 5 nodes of Ranvier and 6 myelinated inter-nodal segments, as myelin is necessary to facilitate the firing of the cell in response to TMS [14][34]. The axonal initiation of the MS-initiated action potential is shown in Supplementary Figure S3. A schematic of the axon morphology is shown in Supplementary Figure S4.

To ensure that the spiking characteristics were in accordance with realistic behaviour for a CA1 pyramidal cell, the axon initial segment had an elevated sodium conductance as well as elevated K_A_ and K_M_ conductances [38]. The values of these conductances were derived by increasing them to a sufficient level where the smaller axon initial segment generated similar electrophysiological properties to the original model with the *HippoUnit* validation battery (see below, [36]).

A list of the specific channels, their conductances, and the passive properties of the different cell compartments is given in in Supplementary Table 1, with properties that have been modified from the original CA1 pyramidal cell model in bold italics. Three properties were ranged within their given regions and the equations governing their values are given in below Supplementary Table 1. None of these ranged variables have been changed from the original model.

### Synaptic models

The excitatory synaptic mechanism used in the plasticity simulations for AMPA/NMDA synapses was the voltage-dependent 4-pathway model of Ebner et al.[30], while for inhibitory synapses the non-plastic Exp2Syn mechanism, with a negative reversal potential of -75 mV was used.

The default parameters for this model were tuned to necortical plasticity data and were not appropriate for a simulation with a larger number of hippocampal synapses. Apart from the parameters in the Ebner model which were derived from direct biophysical measurements, free parameters such as LTP/LTD amplitudes and thresholds were tuned by hand to match the experimental results from Ikegaya et al.[39]. Synaptic weights for the excitatory synapses were uniformly random within a given interval; this interval was manually tuned as for the other parameters. Synaptic weights were chosen with a multiplier between 0.22 and 0.44 (the tuned interval). This was multiplied by the default weight factors of 0.5 for presynaptic weights and 2 for postsynaptic weights. For all simulations except where otherwise noted, 128 excitatory synapses and 18 inhibitory synapses were placed in relative numbers according to Megias (2001)[40]. A table of parameters for the synaptic plasticity model are shown in Supplementary Table 2.

To examine the effect of different synapse locations on the plasticity, we generated a population of different sets of synapse locations for each simulation. This was done by adjusting the seed of the random number generator. The total input synaptic current for each of these models was kept constant; the choice of weights was the same for each model. The same sets of 10 synapse locations were used for all group results shown, to enable direct comparison between the effects of different stimulation protocols.

### Pharmacology *in silico*

To simulate pharmacological perturbations of the model, the following parameter modifications were made to the model. To simulate bicuculline, the weight of all inhibitory synapses was set to zero. To simulate dendritic TTX application, the conductances of all sodium channels in the dendrites of the cell were set to zero. Similarly, to simulate L-type calcium channel block and total calcium channel block, the conductance of these channels in the cell was set to zero.

As the plasticity model is based on NMDA receptor and calcium dependent mechanisms, the NMDA receptor blockade and calcium channel blockade required additional modifications to the synaptic plasticity model itself. For L-type calcium channel blockade, we set the internal variable *N_α_* to zero at each time step, corresponding to the L-type calcium component of presynaptic LTP. NMDA receptor blockade was simulated by setting the parameter *S NMDA*, representing the NMDA fraction of the combined AMPA-NMDA synapse model to zero (thereby removing the synaptic NMDA component), as well as forcing the internal variable *C*, representing NMDA receptor activation, to zero at each time step (thereby disabling the internal synaptic plasticity component associated with NMDA activation). As NMDA blockade would also eliminate the postsynaptic calcium influx, we also set the variable *N_α_* to zero at each time step. To simulate a calcium-free solution, all calcium influx mechanisms were disabled, as well as calcium-dependent variables within the synaptic model.

### Stimulation protocols

When simulating rMS pulses, we also stimulated all Schaffer collateral synapses simultaneously with the application of the rMS pulse to the entire cell. This decision was made based on the assumption that an rMS pulse will cause all (or a large sub-population of) the cells in the target area to fire simultaneously. Given the close proximity of CA1 and CA3, as well as the likelihood that action potentials will be induced in the terminals of axons in the target area without activation of their cell bodies [14], the assumption of near-simultaneous activation of all synapses was implemented in our simulations.To account for the delay in synapse activation, we apply a 1ms delay from the onset of the magnetic stimulus to postsynaptic activation.

In addition to the rMS pulses, we applied a random asynchronous input at 3 Hz to mimic spontaneous neuronal activity in organotypic slice cultures. For the paired local electrical stimulation protocol, a current injection of 3 nA amplitude and 0.5 ms duration was applied to the soma, synchronised with each presynaptic activation.

In the case of the rMS simulations, the TMS waveform generator supplied with *NeMo-TMS* was used to generate a monophasic waveform with an amplitude 275 V/m, which was 60V/m above the minimum required to induce an action potential, which was synchronised with the presynaptic activation of all Schaffer collateral synapses, with this synaptic activation delayed 1ms after the magnetic stimulus to simulate synaptic activation delay.

Likewise, theta burst stimulation simulations utilised a 250 V/m biphasic waveform with synchronous activation of all synapses. The theta burst stimulation protocol consisted of 15 bursts of 5 stimuli at 100 Hz, with each burst separated by 100 ms.

### Validation of the CA1 pyramidal cell model using *HippoUnit* tests

Because the reduced model of the CA1 cell was augmented with an axon and a realistic axon initial segment to enable it to fire in response to TMS pulses, in addition to importing the model into the *NeMo-TMS* framework, re-validation using the *HippoUnit* test battery was required to ensure that our modified model still behaved within acceptable electrophysiological limits for a CA1 pyramidal cell.

All five *HippoUnit* tests [36, 37] were performed - the somatic features test, the back-propagating action potential test, the depolarisation block test, the PSP attenuation test and the oblique integration test. These tests are described in detail in Tomko et al. [36] and no changes were made to the test settings used for validation in that paper. The resulting final error scores are shown in Supplementary Figure S6.

## Results

### A module for simulating TMS-induced long-term synaptic plasticity

Existing modelling tools for TMS integrate anatomically and biophysically detailed neuronal models with electric fields induced by TMS, facilitating the simulation of cellular and sub- cellular voltage and calcium dynamics in response to single and repetitive TMS pulses [14, 34, 41–43]. However, these tools lack the incorporation of experimentally validated plasticity rules necessary for simulating standard LTP induction protocols. To bridge this gap, we have extended the Neuron Modeling for TMS (*NeMo-TMS*) toolbox (Figure 1) with a comprehensive synaptic plasticity model that incorporates a robust four-pathway approach to synaptic modifications, building on our previously established work [30]. This enables the simulation of long-term synaptic changes triggered by local electrical stimulation and rTMS (see below). We integrated this plasticity model with a reduced model of CA1 pyramidal neurons (Figure 1) that has been validated [36] in a comprehensive testing environment called *HippoUnit* [35–37]. The CA1 pyramidal cell model effectively captures a range of neural activites including so- matic spiking, backpropagating action potentials, and dendritic integration of synaptic inputs [36] maintaining a balance between detailed biophysical mechanisms and computational tractability. This integration facilitates the exploration of various LTP/LTD scenarios under rTMS, providing insights into the specific modifications across different anatomical layers and synaptic inputs.

**Figure 1.**
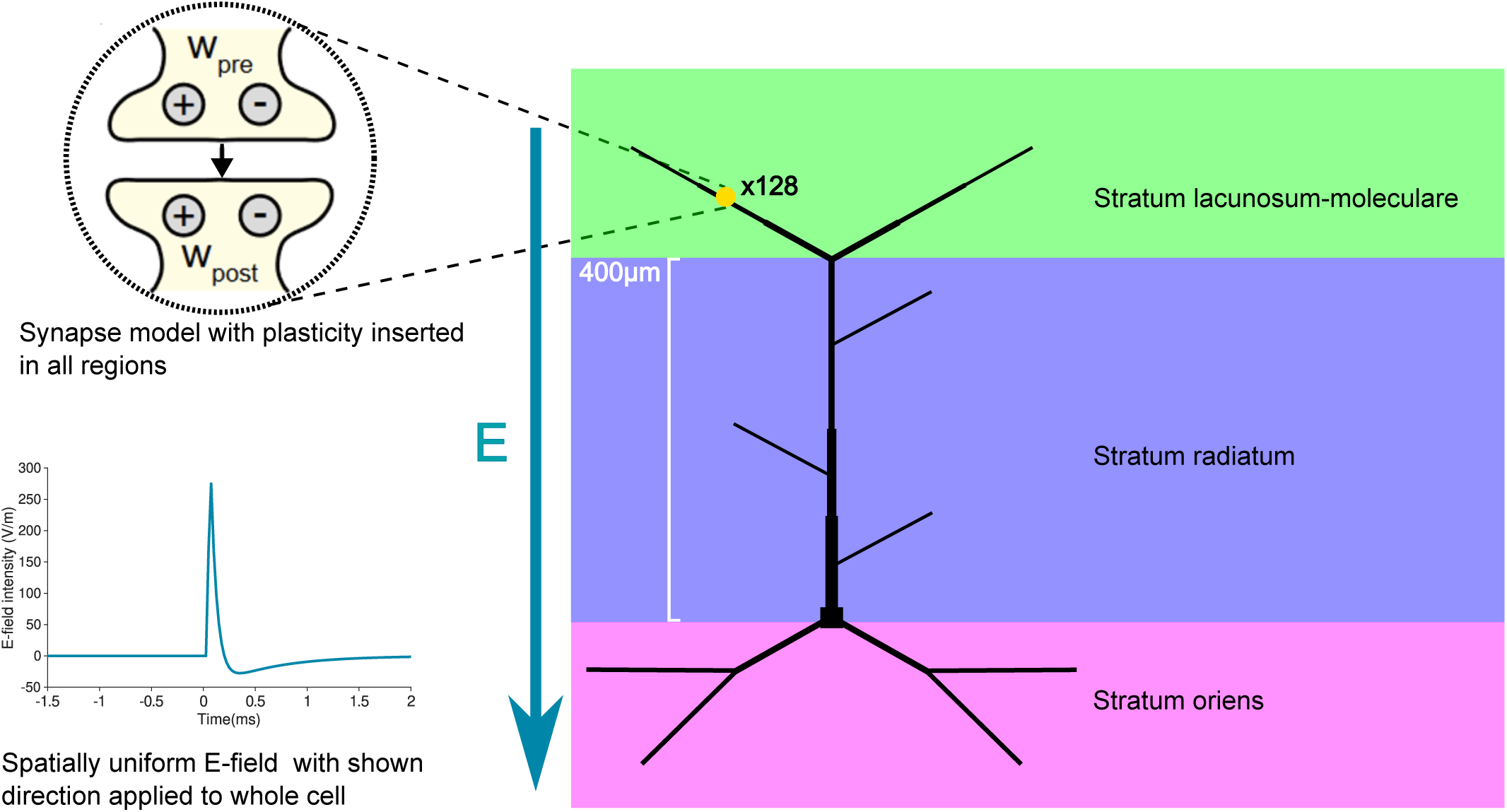
Modelling framework for TMS-induced long-term synaptic plasticity. The Neuron Modelin for TMS (*NeMo-TMS*) toolbox integrates detailed neuronal models with TMS-induced electric fields, allowing the simulation of cellular and subcellular voltage and calcium responses during single and repetitive TM pulses [34]. The direction of the electric field is represented by the vector E. We implement in *NeMo-TMS* a validated model of a CA1 pyramidal cell with detailed biophysics and reduced morphology, capable o generating realistic dendritic and somatic spikes [35, 36]. Relative dendritic diameters are depicted. We introduced a unified voltage-dependent 4-pathway (2xLTP, 2xLTD) model of long-term synaptic plasticity (yellow circle with pre- and postsynaptic LTP (+) and LTD(-)), capable of reproducing the frequency-, timin and location-dependence of synaptic changes [30], into the existing *NeMo-TMS* framework. 128 of plastic excitatory synapses were placed in the morphology.

### Validation of the synaptic plasticity model for LTP/LTD induced by local electrical stimulation

Our main aim was to develop a plasticity modelling framework that would reproduce and explain our previously acquired data on rMS-induced changes of synapses in CA1 pyramidal cells from organotypic slice cultures [31, 44]. However, the unified voltage-dependent model of synaptic plasticity we chose to use was previously tuned to several predominantly neocortical but not hippocampal plasticity datasets [30]. Therefore, we first wanted to test whether the plasticity model could reproduce synaptic plasticity data from classical hippocampal CA3- CA1 synapses. This required tuning of the plasticity model. The parameters used are shown in Table S2.

As a reference hippocampal dataset, we used the frequency-dependent synaptic weight changes from Ikegaya et al. [39], resulting from the induction of LTP or LTD by local extracellular electrical stimulation. In these experiments, the induced LTP/LTD was measured both in the absence of pharmacological perturbation of the culture, and with the application of bicuculline, a GABA_A_ antagonist that suppresses GABAergic inhibition. To simulate the split between control and bicuculline results, we either enabled or disabled the set of inhibitory synapses (see Methods), with the bicuculline case simulated with zero inhibitory synapses enabled. As in Ikegaya et al. [39], the stimulation protocol was a series of 900 pulses delivered to the Schaffer collaterals at a constant frequency of either 1, 10, 30, or 100 Hz. Inhibitory synapses also received an identical stimulus for the simulations where inhibition was active. As in our other simulations, there was an additional 3 Hz Poisson random background activity in all synapses to represent the non-silent nature of slice cultures.

Our model produced results within standard deviation for all conditions except for the 1 Hz control case and the 10 Hz inhibition case (Figure 2). In agreement with the data, significant LTP was only induced above 30 Hz. The change in synaptic weights was negligible at frequencies below 10 Hz, a condition that will be compared later with the rMS results.

**Figure 2.**
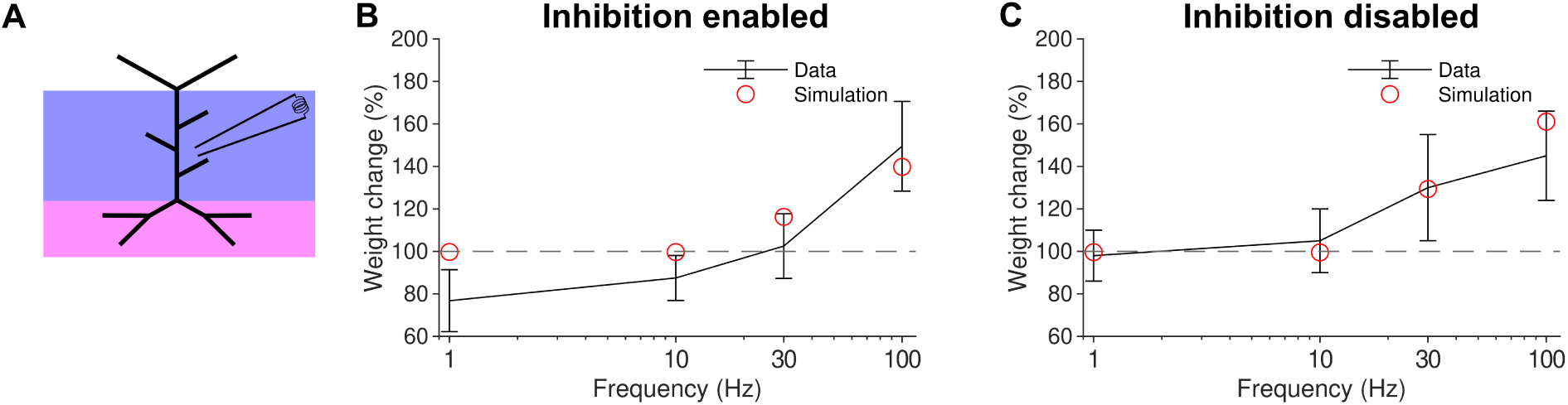
Unifying four-pathway model of long-term synaptic plasticity reproduces frequency dependence of synaptic changes in hippocampal CA1 pyramidal cells evoked by local electrical stimulatio. *(a)* Local electrical stimulation regions (blue purple and magenta) shown on schematic. *(b)* Schaffer collateral weight change for 900 pulses delivered at 1, 10, 30, and 100 Hz with inhibition active to simulate (red control data (blue). *(c)* Same as b but with inhibition disabled to simulate (red) bicuculline data (blue).

These results here indicate that our model can reproduce realistic LTP amplitudes for local electrical stimulation of hippocampal CA1 pyramidal cell synapses. Therefore, the parameters that generated these results (see Table S2) were used for all subsequent simulations.

### Synaptic plasticity model simulates and explains experimentally observed proximal LTP induction by rMS

To determine whether our plasticity model would be applicable to a realistic rMS protocol, we aimed to simulate the plasticity data of Lenz et al. [31]. In this study, an rMS of 900 pulses at 10 Hz was applied to hippocampal slice cultures, inducing strong LTP in the proximal stratum radiatum of CA1. We modelled this with a train of 900 of *NeMo-TMS*’s monophasic pulses with an amplitude set to 275 V/m, clearly above the minimum firing threshold of the CA1 pyramidal cell model. Each *NeMo-TMS* pulse was synchronised with the simultaneous activation of all Schaffer collateral synapses, because we assumed that rMS simultaneously activates not only the CA1 pyramidal cell axon but also the presynaptic axons.

The mean Schaffer collateral synaptic weight increased by 30%, with the LTP induction strongly concentrated in the synapses of the proximal dendrites. This was consistent with an experimental observation of a selective potentiation of proximal synapses in CA1 pyramidal cell dendrites following 10 Hz rMS [31]. Considering only the proximal synapses of the stratum radiatum (defined here as those in the first half of the apical trunk and associated obliques), an average LTP of 65% was achieved, whereas the distal synapses of the stratum radiatum showed no LTP (Figure 3a).

**Figure 3.**
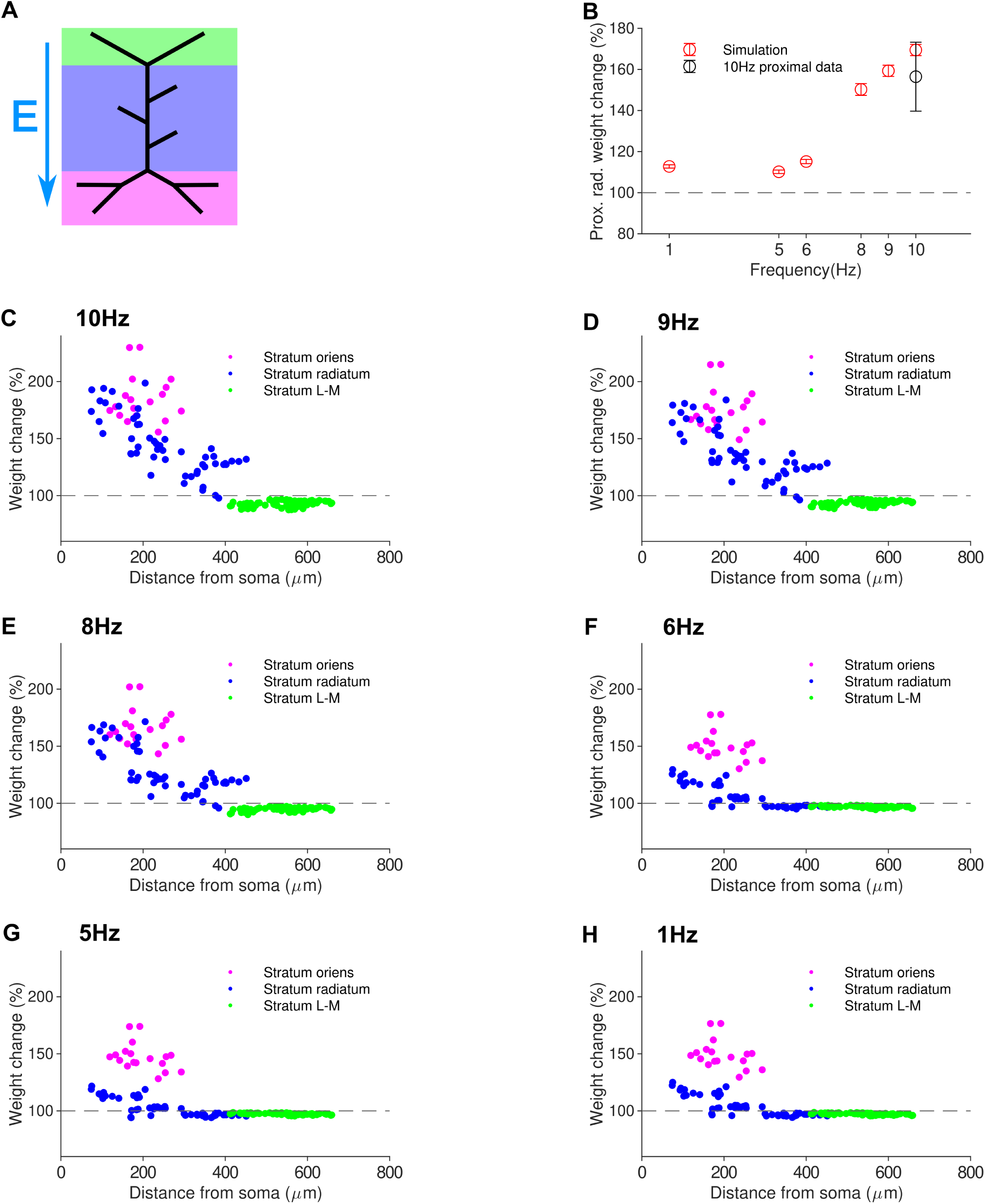
The modelling framework for synaptic plasticity replicates proximal LTP induced by 10 H rMS protocol and predicts increasing LTP with increasing frequency. *a* Schematic of stimulation with electric field vector. *(b)* Mean LTP in the proximal stratum radiatum decreases with decreasing frequenc the induced LTP for 10 Hz rMS is in agreement with experimental data from Lenz (2015). (See next page.) *(c) - (h)* Induction of LTP from 900 pulse rMS protocol at various frequencies. Strong LTP is only seen in the proximal dendrites, with LTP in the stratum radiatum rapidly decreasing with lowe frequency.

To predict the frequency-dependence of synaptic plasticity induced by the rMS protocol, we delivered the stimuli at four additional frequencies of 1, 5, 6, 8, and 9 Hz. Proximal LTP was induced, (Figure 3b), with the amount of potentiation increasing with frequency.

Our simulations here showed that rMS was able to induce significant LTP at much lower frequencies than would be required for purely local electrical presynaptic stimulation ([39], cf. Figure 2). In agreement with experimental data [31], our simulations show that LTP induced by 10 Hz rMS is concentrated in the proximal dendrites, and predict a frequency dependence, with lower frequencies inducing reduced or no LTP.

In additional simulations, biphasic protocol produced less LTP with equivalent stimulation intensity, frequency, and pulse number (Supplementary Figure S1). Increasing stimulation intensity increased LTP amplitude, however these increases became marginal above 300V/m, and no LTP was shown below the cell’s somatic firing threshold of 210V/m. (Supplementary Figure S2).

### Dependence of proximal rMS-induced LTP on dendritic sodium and synaptic NMDA channels

To better understand the mechanistic basis of the experimental and simulation results, we have tested the effects of *in silico* pharmacological perturbations on proximal LTP induced by 10 Hz of 900 rMS (Figure 4). Drugs known to reduce the magnitude of rMS-induced LTP [31] include the sodium channel blocker TTX, L-type calcium channel antagonists and general calcium channel antagonists, as well as NMDA receptor antagonists, since both NMDA channels and voltage-gated calcium channels contribute to overall rMS-evoked LTP [31].

**Figure 4.**
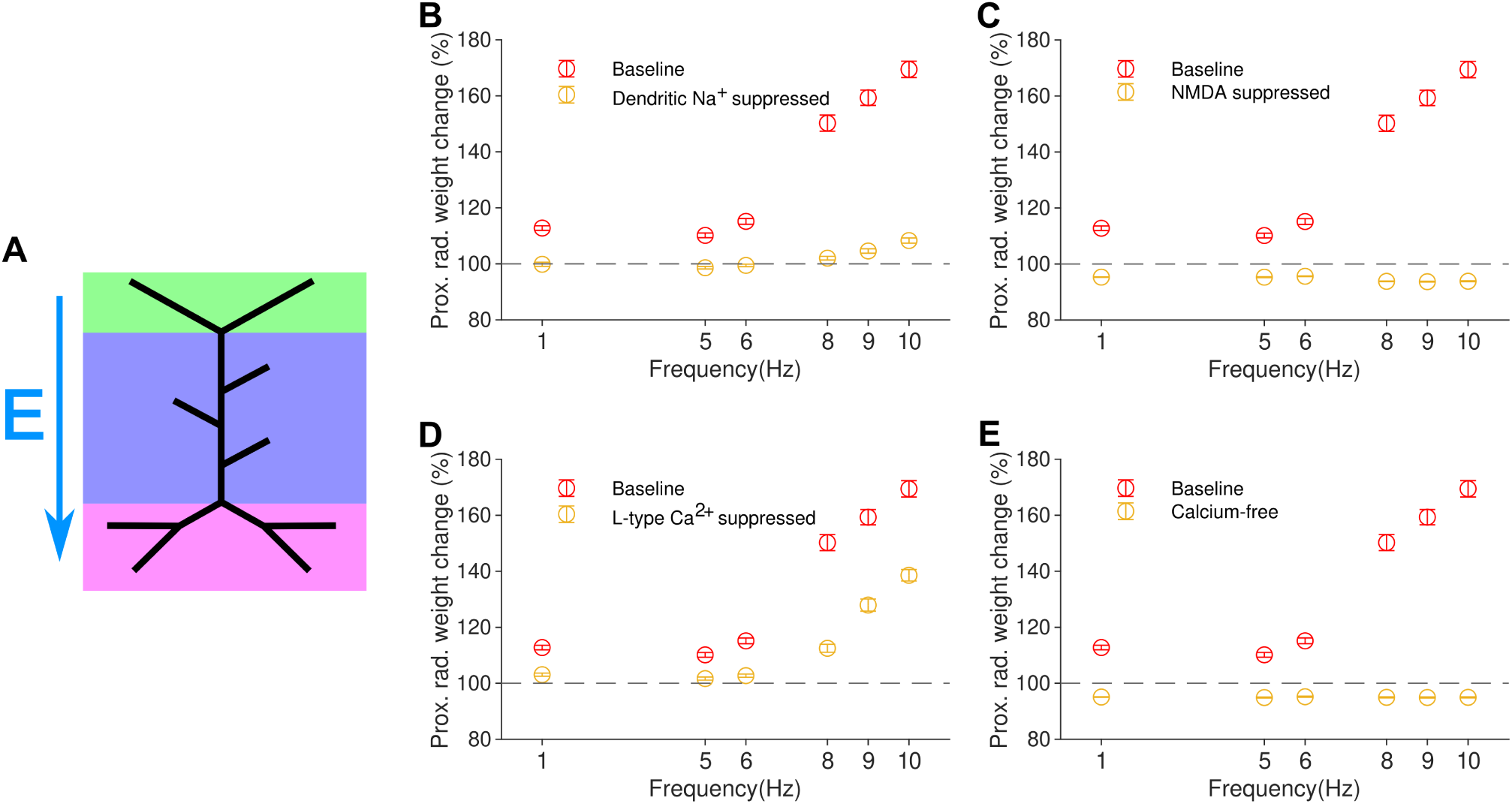
Pharmacology *in silico*: rMS-induced LTP depends on dendritic sodium and calcium, as wel as synaptic NMDA channels. *(a)*: Schematic of electric field for all panels in this figure. Electric field vector direction shown by E; all regions of the cell are synaptically stimulated. *(b)*: Elimination of LTP when dendritic sodium channels are suppressed, simulating TTX application. *(c)*: Elimination of LTP when NMDA-like processes within th synaptic model are disabled *in silico*. *(d)*: Large reduction of LTP induction when L-type calcium channels are suppressed *in silico*. *(e)*: Elimination of LTP observed when calcium-free solution is modelled.

Importantly, in agreement with the experimental data [31], *in silico* suppression (see Methods) of dendritic sodium channels (simulating local TTX application) significantly reduced the amplitude of rMS-induced LTP at all frequencies (Figure 4b). This indicates that our voltage-dependent plasticity rule correctly captures the dependence of proximal rMS-induced synaptic potentiation on sodium channel-mediated depolarisation. In fact, all sodium channels, including somatic ones, were blocked in the experiments of Lenz et al. Our simulations indicate that blocking dendritic sodium channels by local application of TTX would be sufficient to suppress the rMS-induced synaptic potentiation. Similarly, deactivation of NMDA receptors abolished all synaptic plasticity in the simulations (Figure 4c) as observed in experiments [31].

We also tested *in silico* (see Methods) the suppression of L-type calcium channels and the use of a completely calcium-free solution, as these pharmacological perturbations have been reported to reduce rMS-evoked LTP [31]. Suppression of L-type channels caused a large reduction in the amount of induced LTP, although not a complete abolition, as would be expected from Lenz et al. (Figure 4d). A simulated calcium-free solution completely abolished any plasticity (Figure 4e), in agreement with experiments. Taken together, these data-driven pharmacological perturbations *in silico* demonstrate that our modelling framework allows for comprehensive mechanistic simulations of TMS-induced plasticity.

### Simulations predict distal LTP for theta burst stimulation protocols

Our model has so far shown that 10 Hz rMS produces LTP concentrated in proximal dendritic locations. We wondered under what conditions LTP could be induced in the distal dendrites, particularly in the apical tuft of CA1 pyramidal cells. Specifically, we investigated whether it was possible to induce LTP in the apical tuft, where synapses are typically targeted by axons originating in the perforant path from the entorhinal cortex.

The induction of LTP has been shown to require the presence of dendritic spikes [45], mediated by active sodium channels in the dendrites. Accordingly, the unifying plasticity model of Ebner et al. [30] is capable of generating distal LTP in response to local voltage events that cross local plasticity thresholds but do not elicit dendritic and/or somatic spikes. Following its integration into *NeMo-TMS*, we investigated whether similar local synaptic plasticity outcomes could be produced by rMS.

First, to replicate the experiments of Kim et al.[45], we applied their local electrical stimulation protocol of 15 bursts of 5 pulses, with each burst delivered at 100 Hz. In these simulations, no inhibition was present because, as in the experiments, to observe LTP in response to perforant path theta burst stimulation, GABA-A receptors had to be blocked pharmacologically [45]. We also neglected the inter-train interval present in the protocols of Kim et al. because our test simulations showed that inter-train intervals had a negligible effect on synaptic plasticity in this model (as we would expect in a model lacking metaplasticity, where inter-train intervals are longer than the time required for the synaptic model to reset), and would only increase the computational time required. In Kim et al.’s experiments, the soma was prevented from spiking; in our simulations, we found that the somatic voltage deflections from the TBS were very small - on the order of 2 mV, which never produced a somatic spike. As a comparison to these local, purely electrical stimuli, we investigated the effect of rMS delivered in a theta-burst fashion. In these cases, the MS-like stimulus was a biphasic pulse delivered at 275 V/m. To account for the fact that rMS would stimulate many presynaptic neurons, synapses in all layers were activated (as in the 10 Hz rMS simulations above), not just those in the perforant path.

We compared the LTP/LTD distance profiles of all four cases - local electrical TBS with and without inhibition, and rMS TBS with and without inhibition (Fig. 5). With local purely electrical TBS and no inhibition, LTP was observed in the distal tuft, concentrated at the tips (Fig. 5b). LTP was greatly reduced in the presence of inhibition (Fig. 5a). In contrast to local electrical TBS, we observed LTP not only in proximal branches but also in the distal tuft under an MS-TBS protocol, both in the presence and absence of inhibition (Fig. 5c, 5d). Similar to the 10 Hz rMS protocol, LTP induction was also observed in more proximal synapses, to a similar extent as in the distal tuft. In summary, our simulations reproduce and predict the distal strengthening of synapses by electric and rMS TBS, respectively.

**Figure 5.**
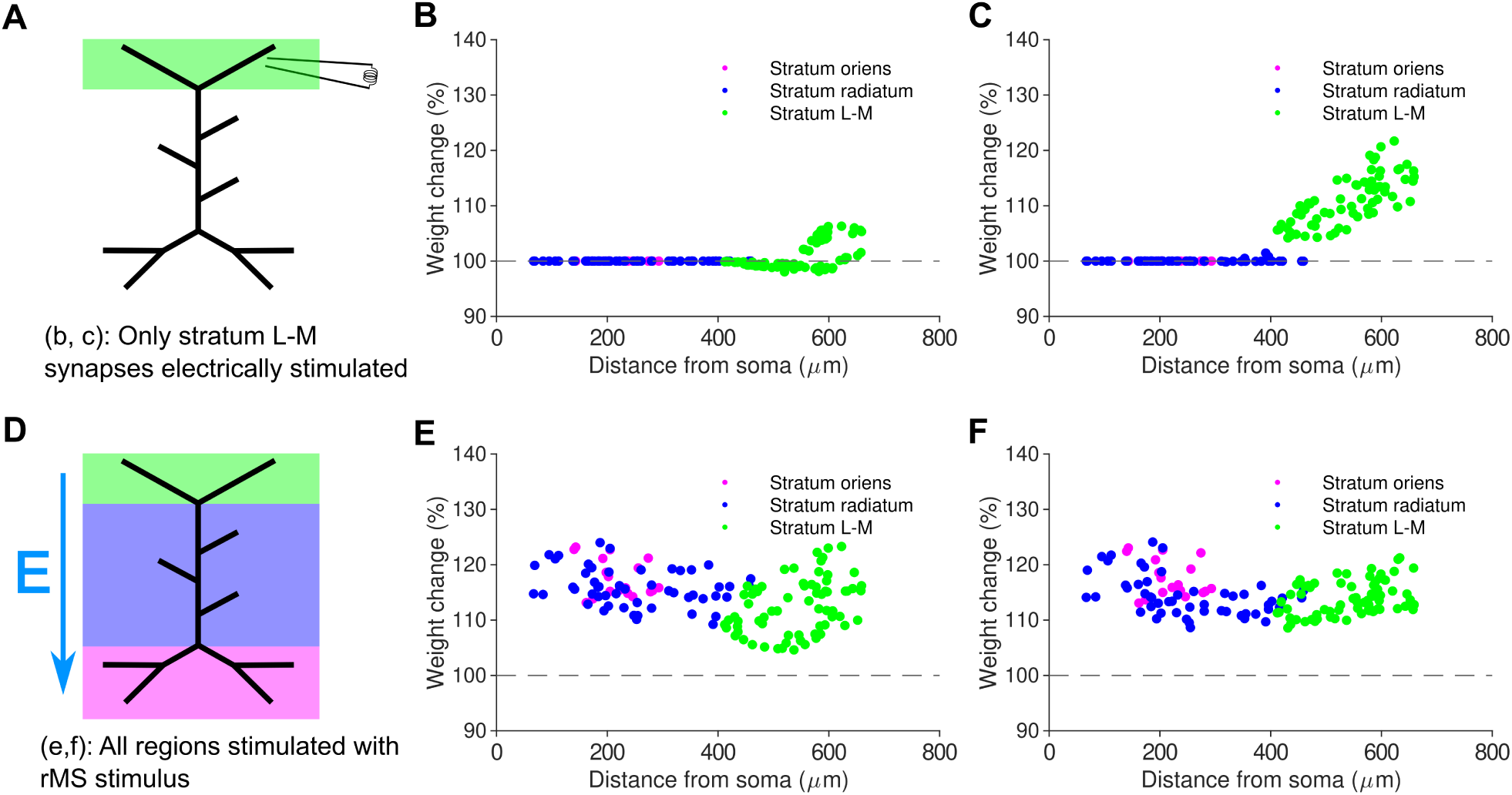
The synaptic plasticity model reproduces distal LTP for local electric TBS and predicts prox mal and distal LTP for rMS TBS. Induction of LTP by local electrical TBS to the perforant path. *(a)*: Schematic of local electric stimulation for panels b and c. L-M: lacunosum-moleculare (perforant path target). *(b)*: Negligible plasticity observed whe inhibition is present. *(c)*: Distal LTP observed when inhibition is absent. *(d)*: Schematic of rMS simulation with electric field vector for panels e and f. *(e)*: LTP in both distal and proximal synapses when rMS is use Unlike local electrical TBS, rMS TBS is able to induce large LTP when inhibition is present. *(f)*: LTP in bot distal and proximal synapses when rMS is simulated and inhibition is disabled.

We observed the induction of dendritic spikes in the distal tuft when local electrical stimulation was delivered to the perforant path synapses in stratum lacunosum-moleculare. Similarly, we observed large-amplitude voltage depolarizations composed of EPSPs, bAPs and presumably also dendritic spikes, when rMS was delivered to the entire cell (Figures 6b,6d). Inhibition suppressed these dendritic voltage events with local electrical stimulation of the perforant path, but not with rMS (Figures 6f, 6h). These somatic and dendritic voltage traces show that local depolarisations that exceed local plasticity thresholds in the dendrite can mechanistically explain why local electric TBS induces distal LTP, whereas rMS TBS induces proximal and distal LTP that is less sensitive to synaptic inhibition.

**Figure 6.**
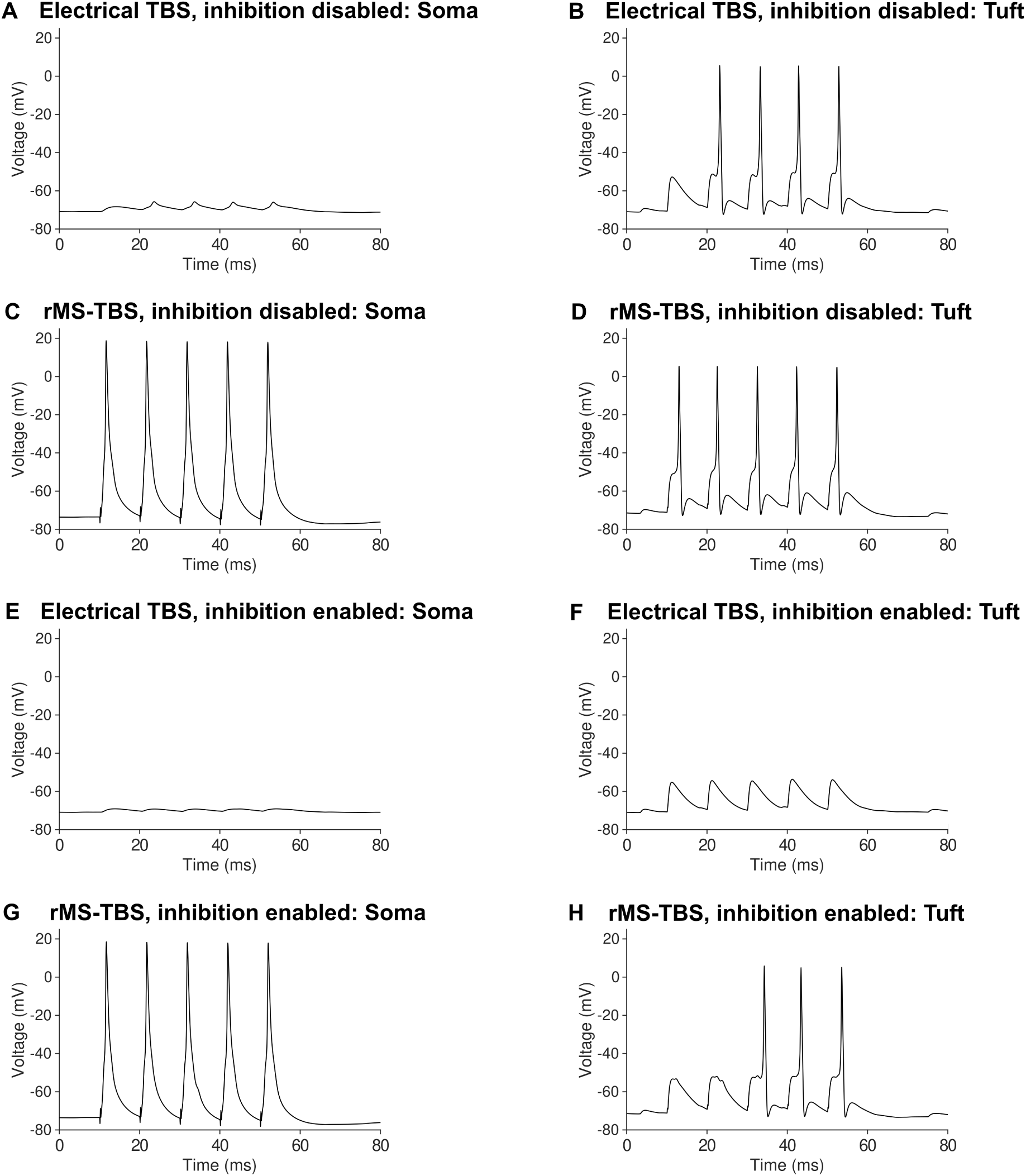
Voltage traces from local electrical and rMS TBS, recorded at the soma and distal apical tuf account for the outcomes of synaptic plasticity. *(a)* 100 Hz local electrical TBS only causes small somatic voltage deflections when inhibition is disabled. *(b* 100 Hz local electrical TBS induces dendritic spikes when inhibition is disabled. *(c)* 100 Hz rMS TBS cause multiple somatic spikes. *(d)* 100 Hz rMS TBS induces dendritic large-amplitude voltage depolarisations when inhibition is disabled. *(e)* 100 Hz local electrical TBS causes almost no somatic voltage deflection when inhibition is enabled. *(f)* 100 Hz local electrical TBS does not induce dendritic spikes when inhibition i enabled. *(g)* 100 Hz rMS TBS causes multiple somatic spikes with inhibition enabled. *(h)* 100 Hz rMS TB induces dendritic large-amplitude depolarisations even when inhibition is enabled.

### Simulated distal LTP evoked by rMS TBS is dependent on sodium den- dritic spikes and less sensitive to inhibition than local electrical TBS

Using local application of TTX, experiments by Kim et al. showed that distal LTP is dependent on sodium dendritic spikes [45]. Our model was able to reproduce this result (Fig. 7) providing further validation of the plasticity model. We eliminated dendritic spikes by simulating an application of TTX to the apical dendrites, which deactivates the sodium channels in the apical dendrites but not in the soma.

**Figure 7.**
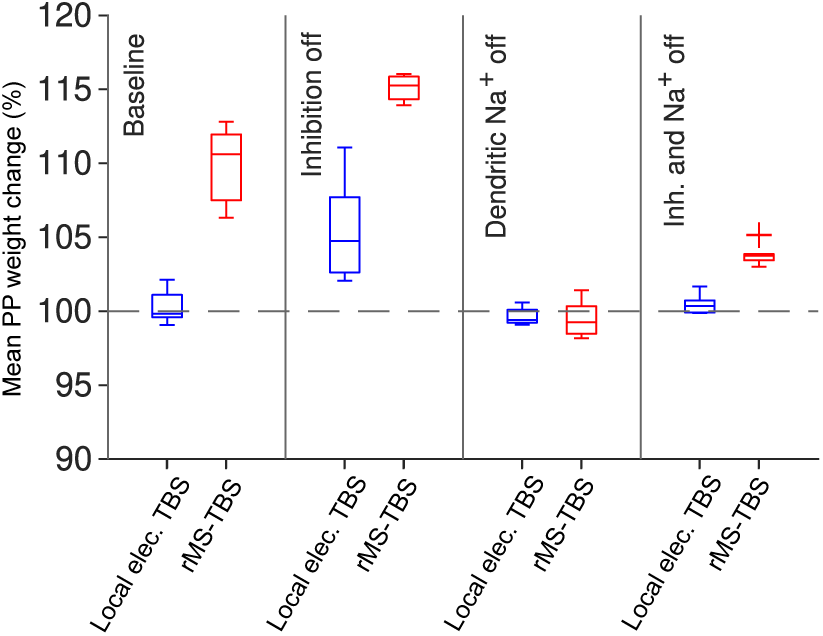
**Distal LTP induced by simulated rMS is sustained by sodium dendritic spikes but less a fected by inhibition than local electrical stimulation**. rMS TBS (red) generates LTP even when local electrical TBS (blue) does not. In the presence of simulated GABAergic inhibition, rMS induces LTP comparable to local electrical stimulation without inhibition (i. bicuculline simulation). When dendritic sodium spikes are blocked (i.e. when local TTX application is simulated), neither of the two stimulation protocols we tested was able to induce LTP in the dendritic tuf Sample of 10 models with different randomised synapse locations.

In the absence of inhibition (bicuculline condition), local electrical TBS was able to restore LTP, whereas rMS TBS induced a slightly greater LTP amplitude as compared to the control condition. In contrast to blocking inhibition, the simulated application of TTX abolished LTP under local electrical TBS and also abolished LTP under rMS TBS.

These results show that our model is capable of producing consistent LTP in the distal tuft of pyramidal cells when local electrical or rMS theta-burst stimuli are applied. This represents a successful reproduction of experimental observations in the case of local electrical TBS. It also predicts powerful LTP when using rMS version of the TBS instead. The model also predicts that this distal LTP will be less diminished by the presence of inhibition in the case of rMS than in the case of local electrical stimulation. However, in both cases, the presence of dendritic sodium spikes is critical for the induction of LTP.

## Discussion

In this work, we have implemented a robust voltage-dependent modelling framework for long-term synaptic plasticity in neurons with extended dendrites. This framework goes beyond point neuron [46] and mean field modelling approaches [47–50] by allowing the simulation of layer and location dependent plasticity effects. Our plasticity simulations in the *NeMo-TMS* toolbox have provided a better mechanistic and quantitative understanding of experimental results on proximal LTP induced by 10 Hz rMS, including pharmacological perturbations using blockers of synaptic and intrinsic ion channels. We made specific predictions for long-term synaptic plasticity induced by lower frequencies of rMS. In addition, we reproduced experimental data on distal LTP induced by local electrical TBS of perforant path synapses and made predictions for proximal and distal LTP induced by rMS TBS, again including pharmacological perturbations. Moreover, our simulations indicate that 1 Hz rMS does not induce long-term depression.

### Modelling TMS-induced synaptic plasticity at subcellular resolution

rMS has previously been shown to induce LTP in a subset of proximal synapses [31]. However, there have been no computational tools to simulate and investigate layer- or compartment-specific rMS-induced synaptic plasticity at a spatio-temporal level with sufficiently high resolution. By incorporating our detailed synaptic plasticity model into the *NeMo-TMS*, it is now possible to simulate TMS-induced plasticity effects at selected spatial and temporal scales. Previously published tools used neural field theory and STDP modelling [51] to predict long-term synaptic changes in populations of neurons driven by repetitive and paired-pulse TMS protocols [47–50]. This type of modelling allowed estimation of the burst rates and number of pulses per burst that result in synaptic weight changes. However, because neurons were not represented as individual cells with dendrites and axons, but as averaged groups of neurons, layer-specific or compartment specific changes of synapses could not be modelled (see also [46]). By combining multi-compartmental neuron modelling with a voltage-based plasticity rule, our work has filled this gap. In addition, while the basic version of *NeMo-TMS* [34] included very simple synaptic modelling (one strong synapse placed in the apical trunk), our current version of *NeMo-TMS* allows arbitrary placement of many synapses, with customisable parameters to determine their distribution within the dendrites of neuronal models. This allows for the investigation of location-dependent dynamics and plasticity of synaptic inputs in neuronal somata and dendrites.

### Modelling distance-dependent gradients of TMS-induced plasticity in neuronal dendrites

In agreement with experiments [31], our simulations showed strongly distance-dependent synaptic changes induced by rMS, with robust LTP in the proximal synapses. Our voltage-dependent plasticity model indicates that proximal LTP can be explained by stronger proximal depolarisation evoked by rMS-evoked backpropagating axonal spikes and their coincidence with proximal postsynaptic responses as compared to distal postsynaptic responses.

Based on the observation of strongly distance-dependent LTP, we asked under what conditions TMS could induce potentiation in distal synapses. We made the prediction that an rMS TBS applied to all synapses and the postsynaptic neuron would generate LTP not only in proximal but also in distal synapses. These predictions can be tested in organotypic hippocampal slice cultures. In these TBS simulations, MS caused the cell to fire in response to almost every stimulus pulse, with little difference with or without inhibition, so the depolarisation at the distal tuft was very similar with or without inhibition. Therefore, because of somatic spike induction, magnetic TBS was able to induce LTP in these distal synapses, unlike local electrical TBS when LTP was blocked by inhibition. However, in the case of TTX simulation (local blockade of dendritic sodium channels), LTP was prevented even with rMS TBS, because inhibition was able to attenuate the backpropagating action potentials from somatic firing, with the backpropagation unable to amplify itself in the absence of sodium channels.

### Assumptions about synaptic activity, magnetic fields and neuronal mor- phology

In this modelling framework, we made a number of physiologically plausible assumptions in both the synaptic model and the presynaptic stimulus. It is known that cells in organotypic slice cultures are not silent [52], so we added a 3 Hz background activity with random jitter to all excitatory and inhibitory synapses. Multielectrode recordings or calcium imaging could be used to better characterise the ongoing neuronal activity in the entorhinal cortex and the CA3 of organotypic slice cultures. Such data could be used to improve the simulations of background synaptic activity.

Similarly, in our simulations, the presynaptic stimulation from both local electric and magnetic stimulation was delivered synchronously to all synapses with a constant delay of 1 ms. In the case of rMS, we believe this is a partially valid assumption because we used a uniform electric field. We assumed that this would have resulted in simultaneous activation of a large proportion of incoming axon terminals. We therefore modelled a simultaneous activation of all synapses, both excitatory and inhibitory. To make these simulations more realistic, random synaptic failures and stochasticity in synaptic delays could be included in the future, based on experimental data.

Furthermore, in our simulations, we imposed the stimulation as a uniform magnetic stimulus. This is a reasonable approximation for simulating neurons in an in vitro environment. However, replicating these results in a more realistic setting including an explicit dish model of the slice culture [53] or a head model and a non-uniform electric field would be an interesting further development due to the spatial heterogeneity and variability of magnetic and electric fields.

It is also worth noting that while our simplified model of the CA1 pyramidal cell closely replicates key dendritic and somatic electrophysiological features of the actual cell [36], it contains significantly less morphological detail than full morphological models of cells. Therefore, applying the plasticity modelling framework to morphologies with dendrites of realistic diameter and branching may improve simulations of localised dendritic voltage-based plasticity [54].

### Metaplasticity, inhibitory plasticity, and clinically relevant stimulation pro- tocols

Future improvements to the *NeMo-TMS* plasticity modelling framework should include metaplasticity, which is observed as persistent shifts in the threshold of LTP/LTD induction that are caused by prior activity of the cell [55]. A modelling implementation of metaplasticity is the BCM (Bienenstock-Cooper-Munro) [56] theory. For stimulation experiments lasting tens of seconds or longer, the combination of a metaplastic model with the current *NeMo-TMS* plasticity model could increase its predictive power. For example, a BCM-like model based on metaplastic STDP [57][58] could be easily integrated with the current four-pathway voltage-dependent plasticity model.

Including metaplasticity would also be useful for simulating the effect of inter-train intervals [47, 50], which we neglect in our TBS simulations. Although originally included in rTMS protocols to avoid overheating the device, they have some effect on plasticity induction [59]. Metaplasticity would also be particularly relevant for investigating the ”priming” of inhibitory theta burst stimulation by repetitive magnetic stimulation, where a low-frequency magnetic stimulus is used to prepare larger magnitudes of LTP induction with a subsequent potentiating stimulus.[60]

Furthermore, given that the presence or absence of inhibition has a modulatory effect on long-term synaptic plasticity [61–63], a more detailed modelling of diverse inhibitory mechanisms would be welcome. Here, we simulated inhibition as being activated simultaneously with excitatory inputs, but simulating feed-forward and feedback inhibition would be more realistic, as the E-I balance in CA1 pyramidal cells depends on complex interactions between different inhibitory cells [64–67].

As we used non-plastic inhibitory synapses, we also neglected possible contributions of inhibitory plasticity [68–71]. It has been observed in the hippocampus that rMS is able to reduce dendritic but not somatic synaptic inhibition [72]. *NeMo-TMS* may be particularly well suited to simulate such compartment-specific changes of GABAergic synapses. Long-term potentiation facilitated by disinhibition from local interneurons is a topic of active research [73], so a model of rMS that releases the inhibitory brake on excitatory plasticity [74] would be desirable. Importantly, inhibitory plasticity is capable of changing the sign of excitatory plasticity [68] and affecting location-specific E-I balance [75]).

With the addition of metaplasticity, inhibitory plasticity, or both, biologically realistic modelling of more complicated clinically relevant stimulation protocols such as intermittent theta burst stimulation (iTBS) [76] [77][6] or continuous theta burst stimulation [78] could be implemented (cf. [47, 50, 79]).

## Conclusions

We have developed a new modelling framework for biophysically detailed simulations of TMS-induced synaptic plasticity and validated it in a well-established model of the CA1 pyramidal cell. We find that our framework is able to reproduce the rMS-induced LTP in the CA1 pyramidal cell model at frequencies well below those that would induce LTP in a local electrical stimulation regime. The rMS-induced synaptic plasticity is distance dependent, occurring very strongly in the proximal dendrites, and also frequency dependent, with stronger LTP observed at 10 Hz and 5 Hz compared to 1 Hz. We predict that magnetic TBS induces LTP in both proximal and distal dendrites of CA1 pyramidal cells.

Extending our modelling framework from the single neuron to the large circuit level and from animal to human synaptic data (e.g. [80]) will open new possibilities for predicting clinically relevant plasticity outcomes of rTMS.

## In brief

Here we develop a validated framework designed for simulating long-term synaptic plasticity induced by TMS. This framework offers accurate predictions of layer-specific effects of rTMS- induced synaptic changes and streamlines the screening of rTMS parameters for inducing synaptic plasticity.

## References

1. Mark Hallett. “Transcranial magnetic stimulation: a primer”. In: Neuron 55.2 (2007), pp. 187–199.

2. Jean-Pascal Lefaucheur, André Aleman, Chris Baeken, David H Benninger, Jérôme Brunelin, Vincenzo Di Lazzaro, Saša R Filipović, Christian Grefkes, Alkomiet Hasan, Friedhelm C Hummel, et al. “Evidence-based guidelines on the therapeutic use of repetitive transcranial magnetic stimulation (rTMS): an update (2014–2018)”. In: Clinical Neurophysiology 131.2 (2020), pp. 474–528.

3. Wan-Yu Hsu, Chia-Hsiung Cheng, Kwong-Kum Liao, I-Hui Lee, and Yung-Yang Lin. “Effects of repetitive transcranial magnetic stimulation on motor functions in patients with stroke: a meta-analysis”. In: Stroke 43.7 (2012), pp. 1849–1857.

[4] Bradley N Gaynes, Stacey W Lloyd, Linda Lux, Gerald Gartlehner, Richard A Hansen, Shannon Brode, Daniel E Jonas, Tammeka Swinson Evans, Meera Viswanathan, and Kathleen N Lohr. “Repetitive transcranial magnetic stimulation for treatment-resistant depression: a systematic review and meta-analysis”. In: The Journal of Clinical Psychiatry. 75.5 (2014), pp. 477–489.

5. [5] Megumi Kinjo, Masataka Wada, Shinichiro Nakajima, Sakiko Tsugawa, Tomomi Nakahara, Daniel M Blumberger, Masaru Mimura, and Yoshihiro Noda. “Transcranial magnetic stimulation neurophysiology of patients with major depressive disorder: a systematic review and meta-analysis”. In: Psychological Medicine 51.1 (2021), pp. 1–10.

6. [6] Richard Morriss, Paul M Briley, Lucy Webster, Mohamed Abdelghani, Shaun Barber, Peter Bates, Cassandra Brookes, Beth Hall, Luke Ingram, Micheal Kurkar, et al. “Connectivity-guided intermittent theta burst versus repetitive transcranial magnetic stimulation for treatment-resistant depression: a randomized controlled trial”. In: Nature Medicine 30 (2024), 403–413.

[7] Xin Wang, Zhiqi Mao, Zhipei Ling, and Xinguang Yu. “Repetitive transcranial magnetic stimulation for cognitive impairment in Alzheimer’s disease: a meta-analysis of randomized controlled trials”. In: Journal of Neurology 267 (2020), pp. 791–801.

8. [8] Ying-hui Chou, Patrick T Hickey, Mark Sundman, Allen W Song, and Nan-kuei Chen. “Effects of repetitive transcranial magnetic stimulation on motor symptoms in Parkinson disease: a systematic review and meta-analysis”. In: JAMA Neurology 72.4 (2015), pp. 432– 440.

9. [9] Rasmus Lorentzen, Tuan D Nguyen, Alexander McGirr, Fredrik Hieronymus, and Søren D Østergaard. “The efficacy of transcranial magnetic stimulation (TMS) for negative symptoms in schizophrenia: a systematic review and meta-analysis”. In: Schizophrenia. 8.35 (2022).

10. [10] Eric M Wassermann and Trelawny Zimmermann. “Transcranial magnetic brain stimulation: therapeutic promises and scientific gaps.” In: Pharmacology & Therapeutics 133.1 (2012), pp. 98–107.

11. [11] Ulf Ziemann. “Thirty years of transcranial magnetic stimulation: where do we stand?” In: Experimental Brain Research 235.4 (2017), pp. 973–984.

12. [12] Florian Muller-Dahlhaus and Andreas Vlachos. “Unraveling the cellular and molecular mechanisms of repetitive magnetic stimulation”. In: Frontiers in Molecular Neuroscience. 6.50 (2013).

13. [13] Hartwig R Siebner, Klaus Funke, Aman S Aberra, Andrea Antal, Sven Bestmann, Robert Chen, Joseph Classen, Marco Davare, Vincenzo Di Lazzaro, Peter T Fox, et al. “Transcranial magnetic stimulation of the brain: What is stimulated?–a consensus and critical position paper”. In: Clinical Neurophysiology 140 (2022), pp. 59–97.

[14] Aman S Aberra, Angel V Peterchev, and Warren M Grill. “Biophysically realistic neuron models for simulation of cortical stimulation.” In: Journal of Neural Engineering 15.066023 (2018).

15. [15] Ulf Ziemann. “TMS induced plasticity in human cortex”. In: Reviews in the Neurosciences. 154 (2004), pp. 253–266.

[16] Tao Tan, Jiacun Xie, Zhiqian Tong, Tiaotiao Liu, Xiaojia Chen, and Xin Tian. “Repetitive transcranial magnetic stimulation increases excitability of hippocampal CA1 pyramidal neurons”. In: *Brain Research* 1520 (2013), pp. 23–35.

17. [17] Mari Ogiue-Ikeda, Suguru Kawato, and Shoogo Ueno. “The effect of repetitive transcranial magnetic stimulation on long-term potentiation in rat hippocampus depends on stimulus intensity.” In: Brain Research 993.1–2 (2003), pp. 222–226.

18. [18] Mari Ogiue-Ikeda, Suguru Kawato, and Shoogo Ueno. “The effect of transcranial magnetic stimulation on long-term potentiation in rat hippocampus”. In: IEEE Transactions on Magnetics 39.5 (2003), pp. 3390–3392.

19. [19] Zaghloul Ahmed and Andrzej Wieraszko. “Modulation of learning and hippocampal, neuronal plasticity by repetitive transcranial magnetic stimulation (rTMS)”. In: Bioelectromagnetics 27.4 (2006), pp. 288–294.

20. [20] Tursonjan Tokay, Norman Holl, Timo Kirschstein, Volker Zschorlich, and Rudiger Kö hling. “High-frequency magnetic stimulation induces long-term potentiation in rat hippocampal slices”. In: Neuroscience Letters 461.2 (2009), pp. 150–154.

[21] Fei Wang, Xin Geng, Hua-Ying Tao, and Yan Cheng. “The restoration after repetitive transcranial magnetic stimulation treatment on cognitive ability of vascular dementia rats and its impacts on synaptic plasticity in hippocampal CA1 area”. In: Journal of Molecular Neuroscience 41 (2010), pp. 145–155.

22. [22] Jun Ma, Zhanchi Zhang, Lin Kang, Dandan Geng, Yanyong Wang, Mingwei Wang, and Huixian Cui. “Repetitive transcranial magnetic stimulation (rTMS) influences spatial cognition and modulates hippocampal structural synaptic plasticity in aging mice”. In: Experimental Gerontology 58 (2014), pp. 256–268.

23. [23] Hui-Yun Yang, Yang Liu, Jia-Cun Xie, Nan-Nan Liu, and Xin Tian. “Effects of repetitive transcranial magnetic stimulation on synaptic plasticity and apoptosis in vascular dementia rats”. In: Behavioural Brain Research 281 (2015), pp. 149–155.

24. [24] N Zhang, M Xing, Y Wang, H Tao, and Y Cheng. “Repetitive transcranial magnetic stimulation enhances spatial learning and synaptic plasticity via the VEGF and BDNF– NMDAR pathways in a rat model of vascular dementia”. In: Neuroscience 311 (2015), pp. 284–291.

25. [25] Yingchun Shang, Xin Wang, Xueliang Shang, Hui Zhang, Zhipeng Liu, Tao Yin, and Tao Zhang. “Repetitive transcranial magnetic stimulation effectively facilitates spatial cognition and synaptic plasticity associated with increasing the levels of BDNF and synaptic proteins in Wistar rats”. In: Neurobiology of Learning and Memory 134 (2016), pp. 369–378.

26. [26] Yue Li, Lulu Li, and Weidong Pan. “Repetitive transcranial magnetic stimulation (rTMS) modulates hippocampal structural synaptic plasticity in rats”. In: Physiological Research. 68.1 (2019), pp. 99–105.

[27] Yechiel Levkovitz, Julia Marx, Nimrod Grisaru, and Menahem Segal. “Long-term effects of transcranial magnetic stimulation on hippocampal reactivity to afferent stimulation”. In: Journal of Neuroscience 19.8 (1999), pp. 3198–3203.

[28] Patricio T Huerta and Bruce T Volpe. “Transcranial magnetic stimulation, synaptic plasticity and network oscillations”. In: Journal of Neuroengineering and Rehabilitation 6.7 (2009).

[29] Roman Gersner, Elena Kravetz, Jodie Feil, Gaby Pell, and Abraham Zangen. “Long-term effects of repetitive transcranial magnetic stimulation on markers for neuroplasticity: differential outcomes in anesthetized and awake animals”. In: Journal of Neuroscience. 31.20 (2011), pp. 7521–7526.

30. [30] Christian Ebner, Claudia Clopath, Peter Jedlicka, and Hermann Cuntz. “Unifying long-term plasticity rules for excitatory synapses by modeling dendrites of cortical pyramidal neurons.” In: Cell Reports 29.13 (2019), pp. 4295–4307.

31. [31] Maximilian Lenz, Steffen Platschek, Viola Priesemann, Denise Becker, Laurent M Willems, Ulf Ziemann, Thomas Deller, Florian Müller-Dahlhaus, Peter Jedlicka, and Andreas Vlachos. “Repetitive magnetic stimulation induces plasticity of excitatory post- synapses on proximal dendrites of cultured mouse CA1 pyramidal neurons.” In: Brain Structure and Function 220.6 (2015), pp. 3323–3337.

[32] Amelie Eichler, Dimitrios Kleidonas, Zsolt Turi, Maximilian Fliegauf, Matthias Kirsch, Dietmar Pfeifer, Takahiro Masuda, Marco Prinz, Maximilian Lenz, and Andreas Vlachos. “Microglial cytokines mediate plasticity induced by 10 Hz repetitive magnetic stimulation”. In: Journal of Neuroscience 43.17 (2023), pp. 3042–3060.

33. [33] Michael L Hines and Nicholas T Carnevale. “The NEURON simulation environment”. In: Neural Computation 9. 6 (1997), pp. 1179–1209.

34. [34] Sina Shirinpour, Nicholas Hananeia, James Rosado, Harry Tran, Christos Galanis, Andreas Vlachos, Peter Jedlicka, Gillian Queisser, and Alexander Opitz. “Multi-scale modeling toolbox for single neuron and subcellular activity under Transcranial Magnetic Stimulation”. In: Brain Stimulation 14.6 (2021), pp. 1470–1482.

[35] Matus Tomko, Lubica Benuskova, and Peter Jedlicka. “A voltage-based Event-Timing-Dependent Plasticity rule accounts for LTP subthreshold and suprathreshold for den- dritic spikes in CA1 pyramidal neurons”. In: Journal of Computational Neuroscience 52.2 (2024), pp. 125–131.

36. [36] Matus Tomko, Lubica Benuskova, and Peter Jedlicka. “A new reduced-morphology model for CA1 pyramidal cells and its validation and comparison with other models using HippoUnit”. In: Scientific Reports 11.7615 (2021).

37. [37] Sára Sáray, Christian A Rössert, Shailesh Appukuttan, Rosanna Migliore, Paola Vitale, Carmen A Lupascu, Luca L Bologna, Werner Van Geit, Armando Romani, Andrew P Davison, et al. “HippoUnit: A software tool for the automated testing and systematic comparison of detailed models of hippocampal neurons based on electrophysiological data”. In: PLoS Computational Biology 17.1 (2021), e1008114.

38. [38] Maarten HP Kole, Susanne U Ilschner, Björn M Kampa, Stephen R Williams, Peter C Ruben, and Greg J Stuart. “Action potential generation requires a high sodium channel density in the axon initial segment”. In: Nature Neuroscience 11.2 (2008), pp. 178–186.

[39] Yuji Ikegaya, Yoko Ishizaka, and Norio Matsuki. “BDNF attenuates hippocampal LTD via activation of phospholipase C: implications for a vertical shift in the fre- quency–response curve of synaptic plasticity.” In: European Journal of Neuroscience 16.1 (2002), pp. 145–148.

40. [40] M Megıas, Z S Emri, T F Freund, and A I Gulyas. “Total number and distribution of inhibitory and excitatory synapses on hippocampal CA1 pyramidal cells.” In: Neuroscience. 102.3 (2001), pp. 527–540.

41. [41] Natalie Schaworonkow and Jochen Triesch. “Ongoing brain rhythms shape I-wave properties in a computational model”. In: Brain Stimulation 11.4 (2018), pp. 828–838.

[42] Sergey N Makarov, Gregory M Noetscher, Edward H Burnham, Dung Ngoc Pham, Aung Thu Htet, Lucia Navarro de Lara, Tommi Raij, and Aapo Nummenmaa. “Software toolkit for fast high-resolution TMS modeling”. In: *bioRxiv* (2019). DOI: 10.1101/643346.

43. [43] Axel Thielscher, Andre Antunes, and Guilherme B Saturnino. “Field modeling for transcranial magnetic stimulation: a useful tool to understand the physiological effects of TMS?” In: 2015 37th Annual International Conference of the IEEE Engineering in Medicine and Biology Society (EMBC). IEEE. 2015, pp. 222–225.

[44] Andreas Vlachos, Florian Müller-Dahlhaus, Johannes Rosskopp, Maximilian Lenz, Ulf Ziemann, and Thomas Deller. “Repetitive magnetic stimulation induces functional and structural plasticity of excitatory postsynapses in mouse organotypic hippocampal slice cultures”. In: Journal of Neuroscience 32.48 (2012), pp. 17514–17523.

45. [45] Yujin Kim, Ching-Lung Hsu, Mark S Cembrowski, Brett D Mensh, and Nelson Spruston. “Dendritic sodium spikes are required for long-term potentiation at distal synapses on hippocampal pyramidal neurons.” In: eLife 4 (2015), e06414.

46. [46] Huan Liu, Lei Guo, Youxi Wu, and Guizhi Xu. “Firing activities of hippocampal CA1 neuron model under electromagnetic stimulation”. In: Nonlinear Dynamics 112 (2024), 9515–9530.

[47] PK Fung and PA Robinson. “Neural field theory of synaptic metaplasticity with applications to theta burst stimulation”. In: Journal of Theoretical Biology 340 (2014), pp. 164– 176.

[48] Marcus T Wilson, DP Goodwin, Philip W Brownjohn, Jonathan Shemmell, and John NJ Reynolds. “Numerical modelling of plasticity induced by transcranial magnetic stimulation”. In: Journal of Computational Neuroscience 36 (2014), pp. 499–514.

[49] Marcus T Wilson, Park K Fung, Peter A Robinson, Jonathan Shemmell, and John NJ Reynolds. “Calcium dependent plasticity applied to repetitive transcranial magnetic stimulation with a neural field model”. In: Journal of Computational Neuroscience 41 (2016), pp. 107–125.

50. [50] Marcus T Wilson, Ben D Fulcher, Park K Fung, Peter A Robinson, Alex Fornito, and Nigel C Rogasch. “Biophysical modeling of neural plasticity induced by transcranial magnetic stimulation”. In: Clinical Neurophysiology 129.6 (2018), pp. 1230–1241.

[51] PA Robinson. “Neural field theory of synaptic plasticity”. In: Journal of Theoretical Biology, 285.1 (2011), pp. 156–163.

52. [52] Marina Samoilova, Kirsten Wentlandt, Yana Adamchik, Alexander A Velumian, and Peter L Carlen. “Connexin 43 mimetic peptides inhibit spontaneous epileptiform activity in organotypic hippocampal slice cultures.” In: Experimental Neurology 210.2 (2008), pp. 762–775.

[53] Ivan Alekseichuk, Sina Shirinpour, and Alexander Opitz. Finite element method (FEM) models for translational research in non-invasive brain stimulation. 2020. DOI: 10.5281/zenodo.3857040.

54. [54] Attila Losonczy, Judit K Makara, and Jeffrey C Magee. “Compartmentalized dendritic plasticity and input feature storage in neurons”. In: Nature 452. 7186 (2008), pp. 436–441.

55. [55] Wickliffe C Abraham. “Metaplasticity: tuning synapses and networks for plasticity”. In: Nature Reviews Neuroscience 9. 5 (2008), pp. 387–387.

[56] Elie L Bienenstock, Leon N Cooper, and Paul W Munro. “Theory for the development of neuron selectivity: orientation specificity and binocular interaction in visual cortex”. In: Journal of Neuroscience 2.1 (1982), pp. 32–48.

[57] Lubica Benuskova and Wickliffe C Abraham. “STDP rule endowed with the BCM sliding threshold accounts for hippocampal heterosynaptic plasticity”. In: Journal of Computational Neuroscience 22 (2007), pp. 129–133.

58. [58] Peter Jedlicka, Lubica Benuskova, and Wickliffe C Abraham. “A voltage-based STDP rule combined with fast BCM-like metaplasticity accounts for LTP and concurrent “heterosynaptic” LTD in the dentate gyrus in vivo”. In: PLoS Computational Biology 11.11 (2015), e1004588.

59. [59] Robin FH Cash, Aisha Dar, Jeanette Hui, Leo De Ruiter, Julianne Baarbé, Peter Fettes, Sarah Peters, Paul B Fitzgerald, Jonathan Downar, and Robert Chen. “Influence of inter-train interval on the plastic effects of rTMS”. In: Brain Stimulation 10.3 (2017), pp. 630–636.

60. [60] Gabrielle Todd, Stanley C Flavel, and Michael C Ridding. “Priming theta-burst repetitive transcranial magnetic stimulation with low-and high-frequency stimulation”. In: Experimental Brain Research 195 (2009), pp. 307–315.

[61] Vincent Paille, Elodie Fino, Kai Du, Teresa Morera-Herreras, Sylvie Perez, Jeanette Hellgren Kotaleski, and Laurent Venance. “GABAergic circuits control spike-timing-dependent plasticity”. In: Journal of Neuroscience 33.22 (2013), pp. 9353–9363.

62. [62] Katharina A Wilmes, Henning Sprekeler, and Susanne Schreiber. “Inhibition as a binary switch for excitatory plasticity in pyramidal neurons”. In: PLoS Computational Biology 12. 3 (2016), e1004768.

63. [63] Kathryn M Murphy, Brett R Beston, Philip M Boley, and David G Jones. “Development of human visual cortex: a balance between excitatory and inhibitory plasticity mechanisms”. In: Developmental Psychobiology 46. 3 (2005), pp. 209–221.

64. [64] Guosong Liu. “Local structural balance and functional interaction of excitatory and inhibitory synapses in hippocampal dendrites”. In: Nature Neuroscience 7. 4 (2004), pp. 373– 379.

[65] David Elfant, Balázs Zoltán Pál, Nigel Emptage, and Marco Capogna. “Specific in- hibitory synapses shift the balance from feedforward to feedback inhibition of hippocampal CA1 pyramidal cells”. In: European Journal of Neuroscience 27.1 (2008), pp. 104– 113.

66. [66] Weiguo Yang and Qian-Quan Sun. “Circuit-specific and neuronal subcellular-wide EI balance in cortical pyramidal cells”. In: Scientific Reports 8. 3971 (2018).

67. [67] Aanchal Bhatia, Sahil Moza, and Upinder Singh Bhalla. “Precise excitation-inhibition balance controls gain and timing in the hippocampus”. In: eLife 8 (2019), e43415.

[68] Lang Wang and Arianna Maffei. “Inhibitory plasticity dictates the sign of plasticity at excitatory synapses”. In: Journal of Neuroscience 34.4 (2014), pp. 1083–1093.

69. [69] Maximilian Lenz, Christos Galanis, Florian Müller-Dahlhaus, Alexander Opitz, Corette J Wierenga, Gábor Szabo, Ulf Ziemann, Thomas Deller, Klaus Funke, and Andreas Vlachos. “Repetitive magnetic stimulation induces plasticity of inhibitory synapses”. In: Nature Communications 7.10020 (2016).

70. [70] Guillaume Hennequin, Everton J Agnes, and Tim P Vogels. “Inhibitory plasticity: balance, control, and codependence”. In: Annual Review of Neuroscience 40 (2017), pp. 557–579.

71. [71] Yue Kris Wu, Christoph Miehl, and Julijana Gjorgjieva. “Regulation of circuit organization and function through inhibitory synaptic plasticity”. In: Trends in Neurosciences 45. 12 (2022), pp. 884–898.

72. [72] Andreas Vlachos, Klaus Funke, and Ulf Ziemann. “Assessment and modulation of cortical inhibition using transcranial magnetic stimulation.” In: e-Neuroforum 23. 1 (2017), pp. 9–17.

73. [73] Julien Artinian and Jean-Claude Lacaille. “Disinhibition in learning and memory circuits: new vistas for somatostatin interneurons and long-term synaptic plasticity”. In: Brain Research Bulletin 141 (2018), pp. 20–26.

74. [74] Maximilian Lenz and Andreas Vlachos. “Releasing the cortical brake by non-invasive electromagnetic stimulation? rTMS induces LTD of GABAergic neurotransmission”. In: Frontiers in Neural Circuits 10. 96 (2016).

75. [75] Tim P Vogels, Henning Sprekeler, Friedemann Zenke, Claudia Clopath, and Wulfram Gerstner. “Inhibitory plasticity balances excitation and inhibition in sensory pathways and memory networks”. In: Science 334. 6062 (2011), pp. 1569–1573.

76. [76] Samuel Bulteau, Veronique Sébille, Guillemette Fayet, Veronique Thomas-Ollivier, Thibault Deschamps, Annabelle Bonnin-Rivalland, Edouard Laforgue, Anne Pichot, Pierre Valrivière, Elisabeth Auffray-Calvier, et al. “Efficacy of intermittent Theta Burst Stimulation (iTBS) and 10-Hz high-frequency repetitive transcranial magnetic stimulation (rTMS) in treatment-resistant unipolar depression: study protocol for a randomised controlled trial”. In: Trials 18.17 (2017).

77. [77] Samuel Bulteau, Andrew Laurin, Morgane Pere, Guillemette Fayet, Veronique Thomas- Ollivier, Thibault Deschamps, Elisabeth Auffray-Calvier, Nicolas Bukowski, Jean-Marie Vanelle, Véronique Sébille, et al. “Intermittent theta burst stimulation (iTBS) versus 10 Hz high-frequency repetitive transcranial magnetic stimulation (rTMS) to alleviate treatment-resistant unipolar depression: A randomized controlled trial (THETA-DEP)”. In: Brain Stimulation 15.3 (2022), pp. 870–880.

[78] Andrei V Chistyakov, Bella Kreinin, Sara Marmor, Boris Kaplan, Adel Khatib, Nawaf Darawsheh, Danny Koren, Menashe Zaaroor, and Ehud Klein. “Preliminary assessment of the therapeutic efficacy of continuous theta-burst magnetic stimulation (cTBS) in major depression: a double-blind sham-controlled study”. In: Journal of Affective Disorders 170 (2015), pp. 225–229.

79. [79] Ying-Zu Huang, John C Rothwell, Rou-Shayn Chen, Chin-Song Lu, and Wen-Li Chuang. “The theoretical model of theta burst form of repetitive transcranial magnetic stimulation”. In: Clinical Neurophysiology 122. 5 (2011), pp. 1011–1018.

80. [80] Maximilian Lenz, Pia Kruse, Amelie Eichler, Jakob Straehle, Jürgen Beck, Thomas Deller, and Andreas Vlachos. “All-trans retinoic acid induces synaptic plasticity in human cortical neurons”. In: eLife 10 (2021), e63026.

